# Posterior integration and thalamo-frontotemporal broadcasting are impaired in disorders of consciousness

**DOI:** 10.1101/2021.11.08.467694

**Authors:** Rajanikant Panda, Ane López-González, Matthieu Gilson, Olivia Gosseries, Aurore Thibaut, Gianluca Frasso, Benedetta Cecconi, Anira Escrichs, Gustavo Deco, Steven Laureys, Gorka Zamora-López, Jitka Annen

## Abstract

The study of the brain’s static and dynamical activity is opening a valuable source of assistance for the clinical assessment of patients with disorders of consciousness. For example, glucose uptake and dysfunctional spread of naturalistic and synthetic stimuli has proven useful to characterize hampered consciousness. However, understanding of the mechanisms behind loss of consciousness following brain injury is still missing. Here, we study the propagation of endogenous and *in-silico* exogenous perturbations in patients with disorders of consciousness, based upon directed and causal interactions estimated from resting-state fMRI. We found that patients with disorders of consciousness suffer decreased capacity for neural propagation and responsiveness to events, and that this can be related to glucose metabolism as measured with [^18^F]FDG-PET. In particular, we show that loss of consciousness is related to the malfunctioning of two neural circuits: the posterior cortical regions failing to convey information, in conjunction with reduced broadcasting of information from subcortical, temporal, parietal and frontal regions. These results seed light on the mechanisms behind disorders of consciousness, triangulating network function with basic measures of brain integrity and behavior.

**Highlights:**

1. Propagation of neural events and network responses are disrupted in patients with DoC.
2. Loss of consciousness is related to the malfunctioning of two neural circuits.
3. Posterior cortical regions lack to integrate information in altered consciousness.
4. Breakdown of information broadcasting of subcortical cortical areas in DoC.
5. Loss of network responses in DoC patients is related to glucose metabolism.

## 1. Introduction

Consciousness is a subjective experience. Internally perceived as the personal experience of “*what is it like, to be you*”, the definition of consciousness and its origin are still a matter of scientific and philosophical debates without consensus (Damasio and Meyer, 2009; Nagel, 1974; Tononi, 2004; Tononi et al., 2016). Within the clinical context, however, practitioners treating patients with severe brain injuries and disorders of consciousness (DoC) face the daily reality to help their patients in the best possible manner, regardless of the exact definition of consciousness. For that, it is important to better understand the mechanisms behind pathological loss of consciousness and its recovery, and to count with tangible correlates that accurately assess the state of the patients. The introduction of neuroimaging proxies can thus help improving both diagnosis and decision making (Owen and Coleman, 2008).

Behavioral assessment such as the response to sensory stimuli, pain or simple commands is the first line of action taken at bedside to evaluate patients. From this perspective, it has proven useful to characterize consciousness based upon two components: wakefulness (the level of arousal) and awareness (the content of consciousness) (Demertzi et al., 2015; Laureys, 2005). Patients with severe brain injury can fall into a coma, which is characterized by the absence of both wakefulness and awareness. Patients surviving coma often recover signs of wakefulness, i.e., eye opening, but without manifestation of awareness of the self nor of the environment. Such state is known as the unresponsive wakefulness syndrome (UWS) (Laureys, 2005). Some of these patients gradually regain awareness and progress into the so-called minimally conscious state (MCS), showing a wider range of non-reflexive behaviors such as visual pursuit, localization to pain or response to simple commands, although their ability to functionally communicate remains hampered (Laureys et al., 2004). While behavioral assessment is the gold standard approach for diagnosis of DoC patients, recently the use of glucose PET (i.e., [^18^F]FDG-PET) has proven valuable to enhance the accuracy of the diagnosis further (Thibaut et al., 2021). Along these lines, the value of auxiliary assessments such as neuroimaging proxies are indicated to refine diagnosis, (Giacino et al., 2018; Kondziella et al., 2020) and specially to gain understanding of the mechanisms behind the loss and the recovery of consciousness that might form the foundation for the development of new treatments.

An upcoming approach to assess brain states relies on the analysis of the brain’s dynamical activity. It is well-known that neural activity is characterized by different frequency bands across sleep stages (Armitage, 1995) or cognitive circumstances, and that local field potentials display intercalated epochs of bursting activity followed by silent periods during anesthesia (Silva et al., 2010). Recent studies have shown that loss of consciousness leads to reduced spontaneous neural activity (Wenzel et al., 2019) and that functional connectivity between brain or cortical regions is also significantly reduced (Barttfeld et al., 2014; Demertzi et al., 2015; Thibaut et al., 2021). Moreover, the fluctuating patterns of functional connectivity are altered during reduced consciousness, with shorter life-times and more random transitions between the patterns as observed in normal awake (Barttfeld et al., 2014; Demertzi et al., 2019; López-González et al., 2021; Luppi et al., 2019).

Observing how external perturbations propagate through the brain constitutes an indirect window to probe brain dynamics, and thus its mechanism, in different states. For example, natural audio-visual stimuli presented to subjects undergoing general anesthesia or within deep sleep are still processed in the sensory cortices but fail to integrate at the higher level cortical regions (Krom et al., 2020; Portas et al., 2000). Application of artificial perturbations such as transcranial magnetic stimulation triggers a response of the stimulated regions that is comparable across conditions, but a rapid decline in the propagation of the signals is found during deep sleep, anesthesia or patients with DoC (Casali et al., 2013; Massimini et al., 2005). These observations have been successfully employed to classify the level of consciousness both in patients and during anesthesia (Casali et al., 2013). However, as the procedure focuses on the description of the whole-brain responses by a single number – the perturbational complexity index – it misses the directionality of the evoked causal interactions. These causal interactions have been demonstrated to be sensitive to different states of consciousness and moreover to hold explanatory power with respect to their neural mechanisms (Seth et al., 2011; Signorelli et al., 2021).

In the present paper, we investigate the capacity of both endogenous and exogenous events to propagate along the brain in patients with DoC as compared to normal wakefulness. By use of model-free and model-based analysis methods, all relevant information to characterize the potential of stimuli to propagate is extracted from the resting-state activity, as measured via functional MRI. Thus, bypassing the need to carry out clinical stimulation protocols. First, we studied how spontaneous endogenous events observed within the resting-state blood oxygenation level dependent (BOLD) signals propagate and are subsequently integrated (Deco and Kringelbach, 2017). We found that the autocovariance relaxation times of the BOLD signals exhibit a spatial distribution in healthy controls which was disrupted in the patients, especially in the UWS group, followed by a significantly reduced capacity to integrate endogenous events. Then, we employed a model-based approach to estimate the pair-wise effective connectivity between brain regions (Adhikari et al., 2021; Gilson et al., 2019, 2016). Since effective connectivity captures the directional causal relations, we could simulate the asymmetrical propagation of exogenous perturbations on the network in order to identify feedforward and backward effective pathways, and to recognize changes in the ability of brain areas to ‘broadcast’ or to ‘receive’ information. In particular, we found two well-differentiated subnetworks with altered propagation properties in the patients. The posterior regions of the cortex fail to convey information, while broadcasting of information is reduced in subcortical, temporal, parietal and frontal regions. These results are in line with the decrease in cerebral glucose metabolism as measured with [^18^F]FDG-PET and evidence that the brain activity in patients with prolonged disorders of consciousness lack of the sufficient capacity for the propagation and the subsequent integration of events, which are necessary conditions for conscious perception.

## 2. Materials and methods

### 2.1. Participants

We included subjects with a pathological reduction or loss of consciousness after severe brain injury, so called disorders of consciousness (DoCs), as well as healthy control (HC) volunteers. Written informed consent was obtained from all HC participants and the legal representative of DoC patients for participation in the study. The local ethics committee from the University Hospital of Liège (Belgium) approved the study. This study includes 40 adult DoC patients, of which 26 were diagnosed in the minimally conscious state (MCS) (7 females, age range 23-73 years; mean age ± SD, 41 ± 13 years) and 14 were diagnosed with the unresponsive wakefulness syndrome (UWS) (7 females, age range 20-74 years; mean age ± SD, 49 ± 16 years). Besides 33 age and gender matched healthy controls (HC) (13 females, age range 19-72 years; mean age ± SD, 40 ± 14 years) without premorbid neurological problems were included. The diagnosis of the DoC patients was confirmed through two gold standard approaches. The first is the repeated behavioral assessment using the Coma Recovery Scale-Revised (CRS-R) by trained clinicians and second, using Fluoro-deoxyglucose Positron Emission Tomography (FDG-PET) neuroimaging as an objective test to complement behavioral assessment according to the procedure described by Stender et al (Stender et al., 2014). Patients were behaviorally diagnosed through the best of at least 5 CRS-R assessments evaluating auditory, visual, motor, oromotor function, communication and arousal (Giacino et al., 2004). Patients for whom these two diagnostic approaches disagreed were excluded from further analysis. Disorders of consciousness occur for a variety of reasons (etiology). Among the 40 patients 17 suffered from anoxia causing widespread neural death and 22 of traumatic brain injuries (TBI), that also includes patients with hemorrhagic stroke and cerebral vascular accident (CVA) leading to more focal lesions. Among the patients diagnosed with UWS, 11 suffered anoxia and 3 TBI whereas the MCS group consists of 6 patients with anoxia and 19 with TBI. Patient specific clinical information is presented in Supplementary Table 1.

### 2.2. MRI Data Acquisition

Structural (T1 and Diffusion Weighted Imaging, DWI) and functional MRI (fMRI) data was acquired on a Siemens 3T Trio scanner. The 3D T1-weighted MP-RAGE images (120 transversal slices, repetition time = 2300 ms, voxel size = 1.0 x 1.0 x 1.2 mm^3^, flip angle = 9°, field of view = 256 x 256 mm^2^) were acquired prior the 10 minutes of BOLD fMRI resting state (i.e., task free) acquisition (EPI, gradient echo, volumes = 300, repetition time = 2000 ms, echo time = 30 ms, flip angle = 78°, voxel size = 3 x 3 x 3 mm^3^, field of view = 192×192 mm^2^, 32 transversal slices). Last, diffusion weighted MRI was acquired in 64 directions (b-value =1,000 s/mm^2^, voxel size = 1.8x1.8x3.3 mm^3^, field of view 230x230 mm^2^, repetition time 5,700 ms, echo time 87 ms, 45 transverse slices, 128x128 voxel matrix) preceded by a single unweighted image(b0).

### 2.3. MRI Data Analysis

#### 2.3.1. MRI data preprocessing

Preprocessing was performed using MELODIC (Multivariate Exploratory Linear Optimized Decomposition into Independent Components) version 3.14, which is part of FMRIB’s Software Library (FSL, http://fsl.fmrib.ox.ac.uk/fsl). The preprocessing consisted of the following steps: the first five functional images were discarded to reduce scanner inhomogeneity, motion correction was performed using MCFLIRT, non-brain tissue was removed using BET, intensity was normalized, temporal band-pass filtering with sigma 100 sec was performed, spatial smoothing was applied using a 5mm FWHM Gaussian kernel, rigid-body registration and single-session ICA with automatic dimensionality. Then noise components and lesion-driven artifacts (e.g., head movement, metal, and physiological noise artifacts) were manually regressed out for each subject. Specifically, FSLeyes in Melodic mode was used to identify the single-subject Independent Components (ICs) into “good” for cerebral signal, “bad” for noise or injury-driven artifacts, and “unknown” for ambiguous components. Each component was evaluated based on the spatial map, the time series, and the temporal power spectrum (Griffanti et al., 2017). FIX was applied with default parameters to remove bad and lesion-driven artifacts components (Griffanti et al., 2017). Subsequently, the Shen et al (2015) functional resting state atlas (without cerebellum) was used for parcellation to obtain the BOLD time series of the 214 cortical and subcortical brain areas in each individual’s native EPI space (Finn et al., 2015). The cleaned functional data were co-registered to the T1-weighted structural image using FLIRT. Then, the T1-weighted image was co-registered to the standard MNI space by using FLIRT (12 DOF), and FNIRT. This transformation matrix was inverted and applied to warp the resting-state atlas from MNI space to the single-subject functional data. Finally, the time series for each of the 214 brain areas were extracted using custom-made Matlab scripts using ‘fslmaths’ and ‘fslmeants’.

#### 2.3.2. Structural Connectivity Matrix

We computed an average whole-brain structural connectivity matrix from all healthy participants as described in our previous study (López-González et al., 2021). Briefly, the b0 image in native diffusion space was co-registered to the T1 structural image using FLIRT. Next, the T1 structural image was co-register to the MNI space by using FLIRT and FNIRT. The resulting transformations were inverted and applied to warp the resting-state atlas from MNI space to the native diffusion space using a nearest-neighbor interpolation method. Then, analysis of diffusion images was performed using the FMRIB’s Diffusion Toolbox (FDT) www.fmrib.ox.ac.uk/fsl. The structural connectivity (SC) mask was obtained by averaging the all HC subjects’ SC matrix and applying a threshold of 80% to maintain the top 20% of strongest connections to binarize the SC. This SC mask was used to constrain the functional connectivity matrix for the whole brain EC computation.

#### 2.3.3. Phase-locking matrices

The instantaneous level of pairwise synchronization was calculated by the phase-locking value between two brain regions. First, the BOLD signals were filtered within a narrowband of 0.01-0.09 Hz. Then the instantaneous phases φ*_k_*(*t*) were computed using the Hilbert transform for each BOLD signal individually. This yields the associated analytical signal which represents a narrowband signal s(t) in the time domain as a rotating vector with an instantaneous phase φ(*t*) and an instantaneous amplitude, A(t). That is, s(t) = A(t)cos(φ(t)). Given the instantaneous phases φ*_j_*(*t*) and φ*_k_*(*t*) calculated two brain regions from their corresponding BOLD signals, the pairwise synchronization *P_jk_* (*t*) was defined as the cosine similarity of the two phases:

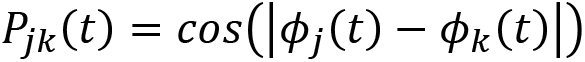

Thus, *P_jk_* (*t*) = 1 when the two regions are in phase, *P_jk_* (*t*) = 0 when they are orthogonal and *P_jk_* (*t*) = -1 when they are in anti-phase (Deco et al., 2017; Ponce-Alvarez et al., 2015).

#### 2.3.4. Intrinsic Ignition

Intrinsic ignition describes the influence of local endogenous events – spontaneously occurring – over the whole-brain network and their subsequent integration (Deco and Kringelbach, 2017). See Deco et al. (2017) for details (Deco and Kringelbach, 2017). Local events are defined as significantly large fluctuations taking place in the resting-state BOLD signals. First, the BOLD signals were transformed into *z*-scores, *Z_i_*(*t*), and binarized by imposing a threshold θ such that the binary signal takes value 1 if *Z_i_*(*t*) > *θ* and 0 otherwise. Here we considered θ = 2 standard deviation (SD). For every endogenous event identified, we calculated the subsequent integration of the event by the network. The integration is assessed using the connectivity out of the phase-locking matrix (Deco et al., 2017; Deco and Kringelbach, 2017). Phase-locking matrices account for the instantaneous level of pairwise synchronization, see below. Integration is then calculated as the area-under-the curve delimited by the size of the largest component in the binarized phase-locking matrix, for all thresholds from 1 to 0 (Adhikari et al., 2017; Deco et al., 2015). The mean intrinsic ignition is finally calculated as the average integration in a time window of 4 TR triggered by all events occurring in the resting-state BOLD for a subject (Deco and Kringelbach, 2017). Higher values of intrinsic ignition correspond to rich and flexible brain dynamics whereas lower values correspond to poor and rigid, structurally driven brain dynamics.

It shall be noted that the analysis of the intrinsic ignition was performed for different thresholds, from 0.5 to 2.5 SD. For ≥2.5 SD some subjects displayed no events in some of the brain regions, therefore we could not analyze the intrinsic ignition for ≥2.5 SD.

#### 2.3.5. Relaxation time constants (τ)

In order to obtain information about the operating regime of brain regions, we measured the relaxation time constant τ from the BOLD signals. Specifically, we measured the time constant of the autocovariance for each brain region individually, using time shifts from 0 to 1 TRs. Given that 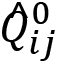 and 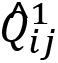 are the zero-lag and 1TR-lag covariance matrices from the empirical BOLD, the time constants τ_*i*_ are calculated as:

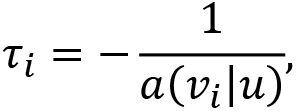

where α (*v*_*i*_|*u*) corresponds to the slope of the linear regression of 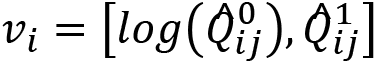 by u=[0,1]. Apart from the information extracted out of the regional time constants, the calculated τ_*i*_ were also employed to inform the estimation of effective connectivity (Gilson et al., 2019).

#### 2.3.6. Estimation of effective connectivity

We estimated whole-brain effective connectivity from the resting-state BOLD signals considering the multivariate Ornstein-Uhlenbeck (MOU) process as the generative dynamical model of the BOLD (Adhikari et al., 2021; Gilson et al., 2016). See Gilson et al (2016) for details (Gilson et al., 2016). The MOU is a model of Gaussian noise diffusion on a network that has been popular to study the relation between the anatomical connectivity and the whole-brain network dynamics (Gilson et al., 2016). Given a structural connectivity matrix **A**, the MOU is defined as:

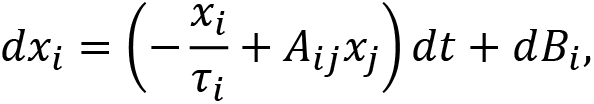

where *x_i_* corresponds to the activity (BOLD signal) of a brain region *i*, τ_*i*_ is the time constant characterizing the exponential decay and dB is a colored noise given by a covariance matrix Σ.

The zero-lag Q^0^ and 1TR-lag Q^1^ covariance matrices of this model can be analytically calculated. The model is thus fitted to empirical data by a Lyapunov optimization procedure such that the distance between the empirical and the estimated Q^0^ and Q^1^ covariances is minimized (Gilson et al., 2016). The optimization process was initialized considering **A** as the binarized structural connectivity matrix in order to restrict the optimization to links identified via diffusion imaging. The estimations were performed using the pyMOU python package.

#### 2.3.7. *In-silico* exogenous perturbational analysis

Considering the MOU as the generative dynamical model for the diffusion of noise in a network, the network responses to local perturbations can be analytically estimated; see Gilson et al. (2019) (Gilson et al., 2019). In particular, we characterize the Green function of the MOU for a given connectivity matrix A. The Green function describes the temporal network responses at times t > 0, due to a unit perturbation applied at a given brain region i at time t = 0. For the MOU process, the spatiotemporal responses are given by:

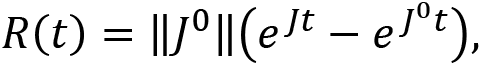

where *J* is the Jaccobian of the MOU process, 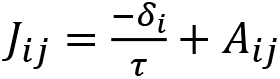 and 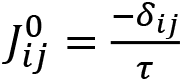 associated to the leakage term alone, characterising the decay rate of the system. ||J^0^|| is a normalization term to make analysis across networks comparable. The response matrices *R*(*t*)encode the spatio-temporal responses to nodal perturbations. In other words, a pair-wise element *R_ij_*(*t*) represents the temporal response of area *j* to a unit perturbation applied on area *i* at time t = 0. This conditional, pair-wise response encompasses all network effects from i to j acting at different time scales. Note that in Ref. Gilson et al., (2019) (Gilson et al., 2019) the network responses *R*(*t*) were referred to as ‘dynamic communicability *C*(*t*) ’. Here we adopted a nomenclature that is clearer and conceptually more precise in order to facilitate the interpretation of results.

As in Ref. Gilson et al. (2019), (Gilson et al., 2019) in the present study, the connectivity matrices **A** are the effective connectivity matrices previously estimated for each subject. Hence, the propagation of responses to exogenous perturbations are constrained by the strength of the directed, causal interactions between every pair of brain regions.

The global network response *r*(*t*) is the sum of all pairwise responses at each time point:

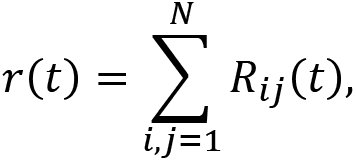

accounting for the total excitability of a network to exogenous perturbation.

Since effective connectivity estimates the directed, causal pairwise interactions between brain regions, and the response matrices *R*(*t*) are constrained upon effective connectivity, *R*(*t*) account for the asymmetric interactions between brain regions. The broadcasting capacity of a region i is calculated as the sum of all responses exerted by region i on the rest of brain areas, and the receiving or integration capacity is given by the sum of responses elicited on region i, by the perturbations at all areas. That is, the broadcasting and receiving capacities of a node are calculated as the row and column sum of the response matrices *R*(*t*) at each time point t:

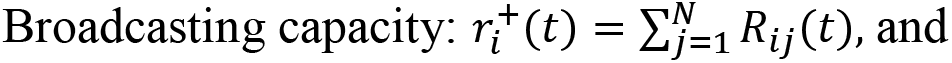

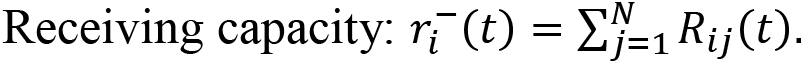

Note that in Refs. Gilson et al. (2019) (Gilson et al., 2019) the broadcasting and receiving capacities are referred to as out-communicability and in-communicability respectively.

#### 2.3.8. Metabolic index using [^18^F]FDG-PET

Alongside the MRI analyses, we have estimated the cerebral glucose metabolism by means of the metabolic index for the best-preserved hemisphere (MIBH) using [^18^F]FDG-PET, that shows high accuracy to discriminate UWS and MCS patients. Behavioral assessment such as the response to sensory stimuli, pain or simple commands is the first line of action taken at bedside to evaluate patients. From this perspective, it has proven useful to characterize consciousness based upon two components: wakefulness (the level of arousal) and awareness (the content of consciousness) (Demertzi et al., 2015; Laureys, 2005). Patients with severe brain injury can fall into a coma, which is characterized by the absence of both wakefulness and awareness. Patients surviving coma often recover signs of wakefulness, i.e., eye opening, but without manifestation of awareness of the self nor of the environment. Such state is known as the unresponsive wakefulness syndrome (UWS). To this end, data acquisition was performed as described elsewhere (e.g., Annen et al (2015) (Annen et al., 2016)). The following steps were followed to calculate MIBH (Stender et al., 2016, 2015). First, we created the patient-specific template for the patients (by taking all DoC patients) and control groups using the procedure described by Phillips and colleagues (Phillips et al., 2011). Then individual images were registered to the appropriate template (for patients or healthy controls) with affine and non-linear registration steps using Advanced Normalization Tools (ANTs version 2.0.3). Images were then segmented into the left cortex, right cortex, extracerebral tissue and normalized based on the metabolism of the extracerebral tissue as described by Stender and colleagues (Stender et al., 2016). Last, metabolic activity was scaled by setting the mean activity of extracerebral regions to an index value of 1 (all values are comprised from 0 to 1). The MIBH was then computed as the highest mean metabolic activity out of the two hemispheres (Stender et al., 2016). We computed differences in the MIBH between UWS and MCS groups using a two-sample t-test. Finally, we correlate the MIBH values with the DoC patients’ network response (i.e., of regions with alterations as compared to healthy controls) using Pearson correlation (considered significant at p<0.05) to explore if the **R**(t) response is grounded by neurobiological underpinnings.

#### 2.3.9. Statistical analyses

For the model free measures, two-sample t-tests were used to assess group differences in global Intrinsic Ignition and relaxation time constant τ at the whole brain level (Bonferroni correction for 3 groups). Specifically, we investigated local between group differences in regional relaxation time constant τ using two-sample t-tests with Bonferroni correction (p<0.05) accounting for the number of regions (i.e., N = 214). Then, we assessed between group differences for the EC links using two-sample t-tests with Bonferroni correction (p<0.05) accounting for 20 % of the number of the structural links between the brain regions (i.e., 9116 links).

For the model based measures, we assessed between group differences in whole-brain total (i.e., receiving and broadcasting) network response. The whole brain network response was calculated as the average network response across all the brain regions. An ANOVA with Tukey post hoc comparison, Bonferroni corrected for 200 timepoints of integration, was employed. Before assessing the local broadcasting and receiving differences, we first evaluated the effect of etiology (anoxia and TBI) versus diagnosis (UWS and MCS) for whole brain early peak responses (i.e., the maximum amplitude of the whole brain network response curve) and late whole brain network responses (i.e., from integration time 60-200 sec (modeled time), based on the findings of the ANOVA described above) using a linear regression model as implemented in MATLAB (i.e., fitlm function). Both the mean effects for diagnosis and etiology were considered as well as their interaction. Regional modulation of communicability was evaluated only for factors with a significant main effect on global communicability.

To investigate local broadcasting and receiving properties, we considered the area under the receiving and broadcasting curves (i.e., from integration time 60-200 sec modeled time) separately for every brain region. We identified regions with relatively high difference in network responses within groups (i.e., HC, UWS and MCS separately) with a one-sample t-test with Bonferroni correction (p<0.05) accounting for the number of regions (i.e., N = 214). We identified the regional dominance for broadcasting and receiving by subtracting the AUCs for the regional receiving from the broadcasting curves and performed a within-group t-test to identify regions with specific functional specialization (considered significant at p<0.05 Bonferroni corrected for N = 214). Last, between group differences in regional receiving and broadcasting information were assessed with two-sample t-tests with Bonferroni correction (p<0.05) accounting for the number of regions (i.e., N = 214).

We computed differences in the MIBH between UWS and MCS groups using a two-sample t-test. Finally, we correlate the MIBH values with the DoC patients’ network response (i.e., of regions with alterations as compared to healthy controls) using Pearson correlation (considered significant at p<0.05) to explore if the computational the network response is grounded by neurobiological underpinnings.

#### 2.3.10. Data Availability

Phase interaction matrices for the fMRI connectivity are available at: https://search.kg.ebrains.eu/instances/Dataset/775c7858-2305-4a56-8bd6-865c4ab5dd4f. After the acceptance of the manuscript, the code used for this study will be available at: https://github.com/RajanikantPanda/Ignition_and_Information_flow_for_DOC

## 3. Results

This study comprises resting-state fMRI (eyes-closed) of 33 healthy control (HC) subjects, 14 patients with unresponsive wakefulness syndrome (UWS) and 26 patients classified as in minimally conscious state (MCS). The diagnoses were made using repeated CRS-R assessments and confirmed with FDG-PET neuroimaging (Stender et al., 2014) to avoid including MCS* patients (Thibaut et al., 2021).

### 3.1. Global integration of local endogenous events is hampered in lower conscious states

We started this study by investigating whether endogenous spontaneous events occurring locally propagate differently depending on the level of consciousness, across healthy controls, MCS or UWS patients. For that, we employed the *intrinsic ignition* measure (Deco and Kringelbach, 2017). The level of global intrinsic ignition for a subject is calculated as the average integration triggered by all endogenous events identified in a resting-state BOLD session, see Methods. As shown in **Fig. 1a**, the mean intrinsic ignition driven by events of 2SD BOLD signal threshold was lowest in UWS patients implying that the endogenous BOLD events lead to a lower network response than in healthy controls and in MCS patients (HC = 0.82±0.02, UWS = 0.78±0.02, MCS = 0.79±0.02, HC vs. UWS t(45)=6.0, p<0.0001, HC vs. MCS t(57)=4.6, p<0.0001, MCS vs. UWS t(38)=2.0, p=0.029). It shall also be noted that the number of observed intrinsic events was lowest in UWS patients, intermediate in MCS patients and highest in healthy controls (HC = 14.1±3.6, UWS = 7.6±2.9, MCS = 11.0±3.6, HC vs. UWS t(45)=5.9, p<0.0001, HC vs. MCS t(57)=3.3, p=0.0017, MCS vs. UWS t(38)=2.9, p=0.0048). We found similar patterns in intrinsic ignition for healthy control and DoC patients for other thresholds (i.e., 0.5, 1.0 and1.5 SD; see **Supplementary Fig. 1**).

**Figure 1.**
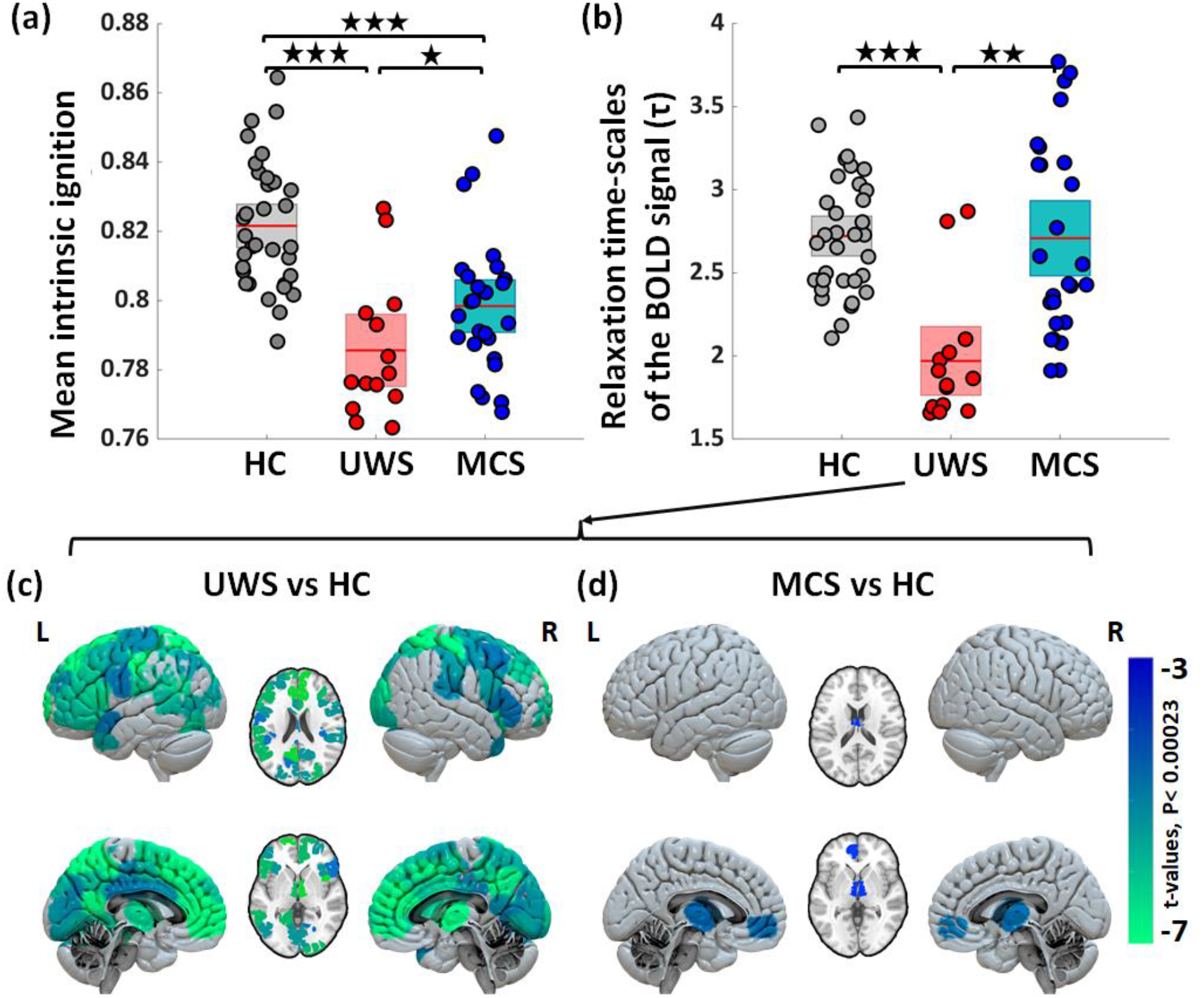
Changes in endogenous properties from resting-state BOLD signals. Healthy controls (HC), unresponsive wakefulness syndrome (UWS) and minimally conscious state (MCS) (a) Comparison of mean intrinsic ignition for the three groups, illustrating the reduced capacity to integrate endogenous spontaneous events in patients with DoC. (b) Relaxation time-scales of the BOLD signals (τ) at the whole-brain level shows significant reductions in UWS and MCS patients compared to HC. Stars reflect the Bonferroni corrected (for three groups) significance levels (*=p-value<0.05; **= p-value<0.001; ***= p-value<0.0001). (c, d) Maps of significant differences in regional distributions of τ between patients and controls. The color bar represents the t-values of significant between-group differences (Bonferroni corrected for 214 tests, p-value=0.05).

### 3.2. Shorter relaxation-time of BOLD signals in low levels of consciousness

Measuring time-scales from signals can reveal changes in the underlying mechanisms controlling the local dynamics and determining their operating regime. Specifically, the autocovariance profile of the BOLD for each brain area estimates the duration for which the signal is altered before going back to pre-event baseline activity (Murray et al., 2014). Here, we measure the autocovariance time constant τ, also called the *relaxation time* or *memory depth* in the literature. Large τ implies a longer lingering effect of a signal after an event or perturbation before it decays, thus suggesting that the brain region might remain available for processing longer.

At a whole-brain level, averaging over the τ_*i*_ for all regions in one subject, we found that τ was shorter in UWS patients (1.96±0.38) than in healthy controls (2.72±0.35; t(45)=6.5, p<0.0001) and MCS patients (2.70±0.58; t(38)=4.2, p<0.001), see **Fig. 1b**. Looking at the region-wise spatial distributions, we found that in healthy controls τ_*i*_ is heterogeneously distributed showing a gradient with shorter time constants in subcortical areas (τ_*i*_ ∼ 1.5 sec) and longer (τ_*i*_ ∼3.5 sec) in the frontal and in the parietal areas, see **Supplementary Fig. 2**. Importantly, the diversity of relaxation times is lost in the UWS patients with τ_*i*_ being homogeneously distributed and dominated by small values. Compared to healthy controls, the decrease of τ_*i*_ in UWS patients was most predominant in the bilateral thalamus, right caudate, left hippocampus, parahippocampus, bilateral posterior, middle and anterior cingulate, insula, inferior, middle, superior and dorsolateral frontal areas, **Fig. 1c** **and Supplementary Table 2**. In the case of MCS patients the heterogeneity of τ_*i*_ distribution was practically recovered, **Supplementary Fig. 2**. Compared to the healthy controls, in MCS patients τ_*i*_was lower only in the bilateral thalamus and left medial prefrontal cortex, see **Fig. 1d** and Supplementary Table 2.

So far, the results obtained for the intrinsic ignition and the distribution of relaxation time constants indicate a breakdown in the spatiotemporal structure of the BOLD signals that involves reduced propagation and integration capabilities of endogenous events in DoC patients, especially in the UWS group. For the remaining of the paper we shift to model-based analyses.

### 3.3. Whole-brain effective connectivity shows altered causal interactions in DoC patients

In order to identify alterations to the causal relations between the brain regions, we estimated whole-brain effective connectivity (EC) from the resting-state BOLD for each subject. The estimation of EC considers a model of Gaussian noise diffusion – the multivariate Ornstein-Uhlenbeck – on top of the anatomical connectivity as the generative dynamics (Adhikari et al., 2021; Gilson et al., 2016) in order to capture the origin of the fluctuations in the BOLD; see **Fig. 2a** and Methods for further details. In short, EC estimation consists of identifying the most likely causal interactions that give rise to the observed BOLD signals, fitting both the interaction strengths between all pairs of ROIs and the levels of noise to stimulate each ROI, **Fig. 2a**.

**Figure 2.**
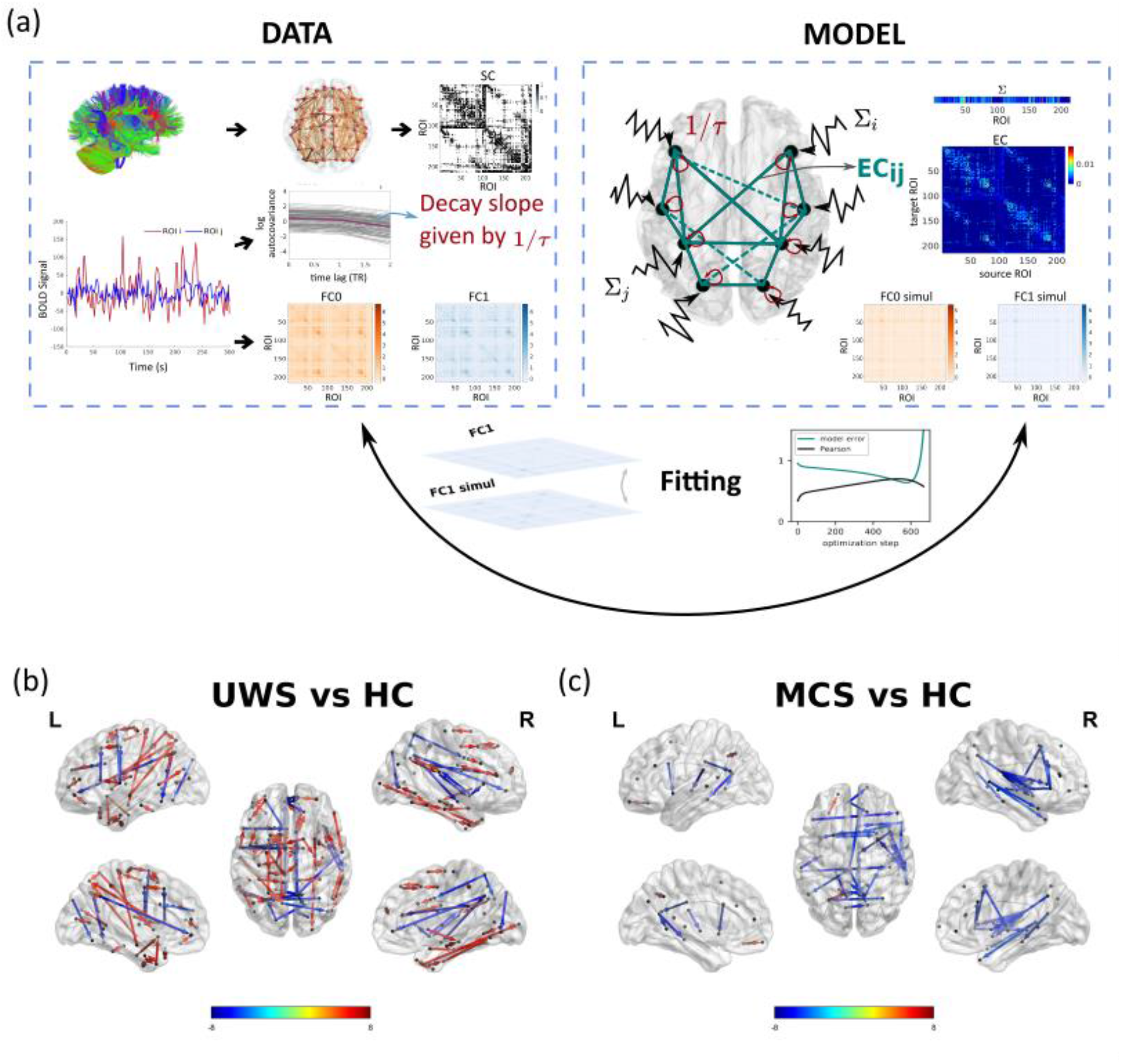
Comparison of effective connectivity (EC) between healthy controls and patients. (a) Schematic representation of the fitting procedure leading to estimation of EC. Considering a model of noise diffusion – the multivariate Ornstein-Uhlenbeck process – the whole-brain network model is constrained using structural connectivity obtained from diffusion imaging and then fitted to reproduce the empirical resting-state data. In particular, to fit the zero-lag and 1TR-lag covariance matrices (FC0 and FC1), and the regional noise level Σi. (b, c) Maps of significantly different EC connections between patients and controls. UWS patients show connections with both decreased and increased EC (decreased in fronto-temporal, frontal-parietal and midline regions; increased in subcortical and wide cortical areas). Blue and red arrows indicate lower and higher EC respectively in patients as compared to HC subjects. MCS patients show decreased EC in fronto-temporal and interhemispheric midline connections. The directional connections in the glass brain represents connections with significant between-group differences (Bonferroni corrected for 11395 tests, p-value < 0.05) are represented.

At the whole-brain level, averaging across all EC links, we found that the EC of the UWS patients was higher than for the healthy controls or in MCS patients (HC = 0.015±0.002, UWS = 0.019±0.004, MCS = 0.014±0.003; HC vs. UWS t(45)=-4.7, p<0.0001, MCS vs. UWS t(38)=-4.0, p<0.001). A closer inspection of the pair-wise EC values revealed the presence of links that either increased or decreased in the UWS patients in respect to the healthy controls, **Fig. 2b**. The UWS patients showed increased EC for connections between subcortical and cortical regions (thalamus, caudate and putamen), but decreased EC in connections spanning posterior (i.e., parietal, occipital) to frontal (i.e., temporal and frontal) regions as well as between midline posterior regions (parietal, occipital) and middle frontal regions. The MCS patients showed especially lower EC in interactions from posterior to frontal and temporal regions and midline regions encompassing the middle prefrontal and posterior cortex and the thalamus, see **Fig. 2c**, including regions important for long range connectivity and overlapping with key areas of the Default Mode Network.

### 3.4. Altered spatiotemporal propagation of exogenous perturbations

Having identified changes in specific pair-wise EC connections for both UWS and MCS patients, the question is now how do those alterations affect the propagation of information in the brain. To answer this question, we perform an *in-silico* perturbational study to assess how exogenous perturbations, applied to individual ROIs, spread along the network. Considering the same generative dynamical model as for the EC estimation, the effect of regional perturbations on the rest of the network can be analytically estimated, (Gilson et al., 2019) see Methods. The spatiotemporal responses of nodal perturbations are encoded into the temporal response matrices R(t). The evolution of response matrices for the three study cases are shown in **Fig. 3a**. Specifically, a pair-wise element Rij(t) represents the temporal response of area j to a unit perturbation applied on area i at time t = 0. This conditional, pair-wise response encompasses all network effects from i to j acting at different time scales.

**Figure 3.**
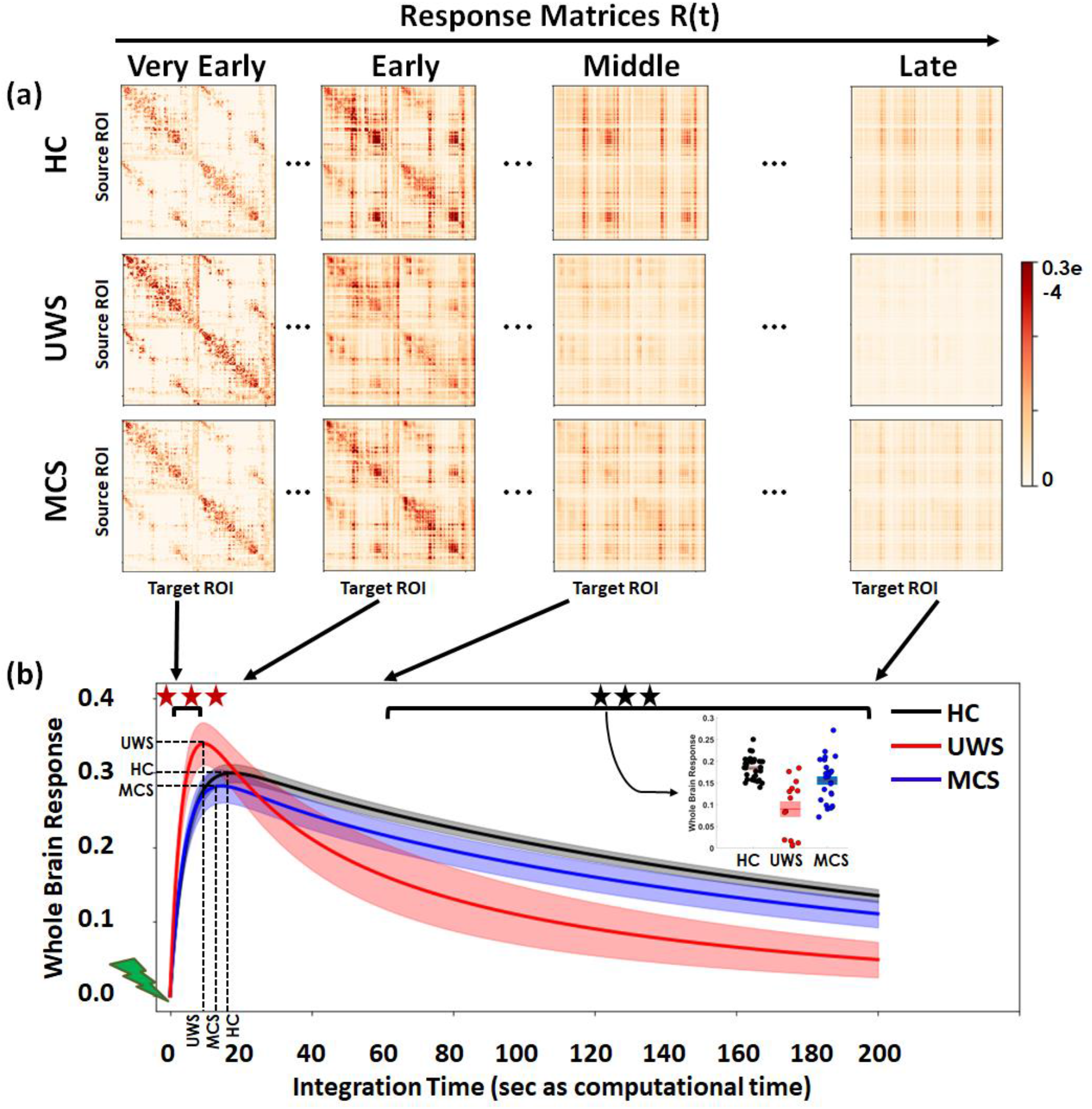
*In-silico* propagation of exogenous perturbations. (a) Temporal evolution of the response matrices R(t) for healthy controls (HC) and patients (unresponsive wakefulness syndrome, UWS; minimally conscious state, MCS) at different times (early = 2 sec, middle = 20 sec, late 60 sec and very late = 200 sec). Matrix elements Rij(t) represent the conditional response at region j due to a unit perturbation applied at region i at time t = 0. Note that here time corresponds to the arbitrary simulation time after the *in-silico* perturbation is applied and thus it does not correspond to actual time, although the time-constants governing the evolution were estimated from the BOLD signals. The colorbar represents the relative strength of the response between brain regions (unitless). (b) Whole brain response curves for the three study cases reflecting the sum of all pair-wise responses at each time point post-stimulus. Shaded areas represent the 95% confidence intervals across subjects. Black stars indicate a difference in global responses between all three groups (Bonferroni corrected for 100 tests/time points, p-value=0.05). Red stars indicate the early epoch during which UWS patients display a larger response than HC and MCS (Bonferroni corrected for 100 tests/time points, p-value=0.05). Inset: Area-under-the-curve for the three global response curves in the time range t = 60 – 200 sec, quantifying the differences across the three groups.

**Figure 3a** illustrates how the patterns of responses are progressively reshaped over time for the three study groups – healthy controls, UWS patients and MCS patients. The global brain responses (sum over all pair-wise responses) are shown in **Fig. 3b**. As seen, the global responses undergo a transient peak short after the initial perturbations and then decay as the effects of the stimuli dilute with time and the system relaxes back to its stationary state. This relaxation is also observed by the homogenization of the response matrices at the longer latencies in **Fig. 3a**. The global response curves for controls and MCS groups follow quite a similar behavior, both peaking at 18.2±2.9 and 15.6± 3.7 seconds respectively and taking peak values 0.30±0.03 and 0.28± 0.05. In the UWS patients, however, the global response peaks sooner (10.6±2.9 sec) (HC vs. UWS t(45)=8.2, p<0.0001, HC vs. MCS t(57)=3.0, p=0.0036, MCS vs. UWS t(38)=4.3, p < 0.001) and displays a higher peak (0.34±0.05) (HC vs. UWS t(45)=-3.3, p= 0.0019, MCS vs. UWS t(38)=-3.1, p = 0.0031) than for the controls and MCS groups, but then it decays notably faster. Quantitatively, we found that the area-under-the-curve in the time spanning 60-200 sec (modeled time) significantly decreases for the UWS group (0.08±0.06) and MCS patients (0.15± 0.04) compared to healthy controls (0.18±0.02) (HC vs. UWS t(45)=7.1, p<0.0001, HC vs. MCS t(57)=2.9, p=0.005, MCS vs. UWS t(38)=3.6, p < 0.001).

### 3.5. Broadcasting and integrative capabilities of brain regions across states of consciousness

We then explored the response of single regions within the network by column- and row-wise exploration of the response matrices **R(t)** in **Fig. 4a**. Since EC identifies the directed causal interactions, this allows us to study the input and output relations for each area in respect to the exogenous perturbations. The row sum of the response matrices **R**(t) represent the broadcasting capacity of a region (i.e., the response that a perturbation in one region elicits on all other areas) and the columns describe the integrative capacity of the brain region (i.e., how much is a region affected by the perturbations applied to all other areas) (Gilson et al., 2019). The temporal evolution of broadcasting and receiving of single regions shows distinct functional roles for various regions in healthy controls. This functional specialization is much reduced in MCS patients, and almost absent in UWS patients, in which the regional roles within the response to in-silico perturbation in the network is homogenized.

**Figure 4.**
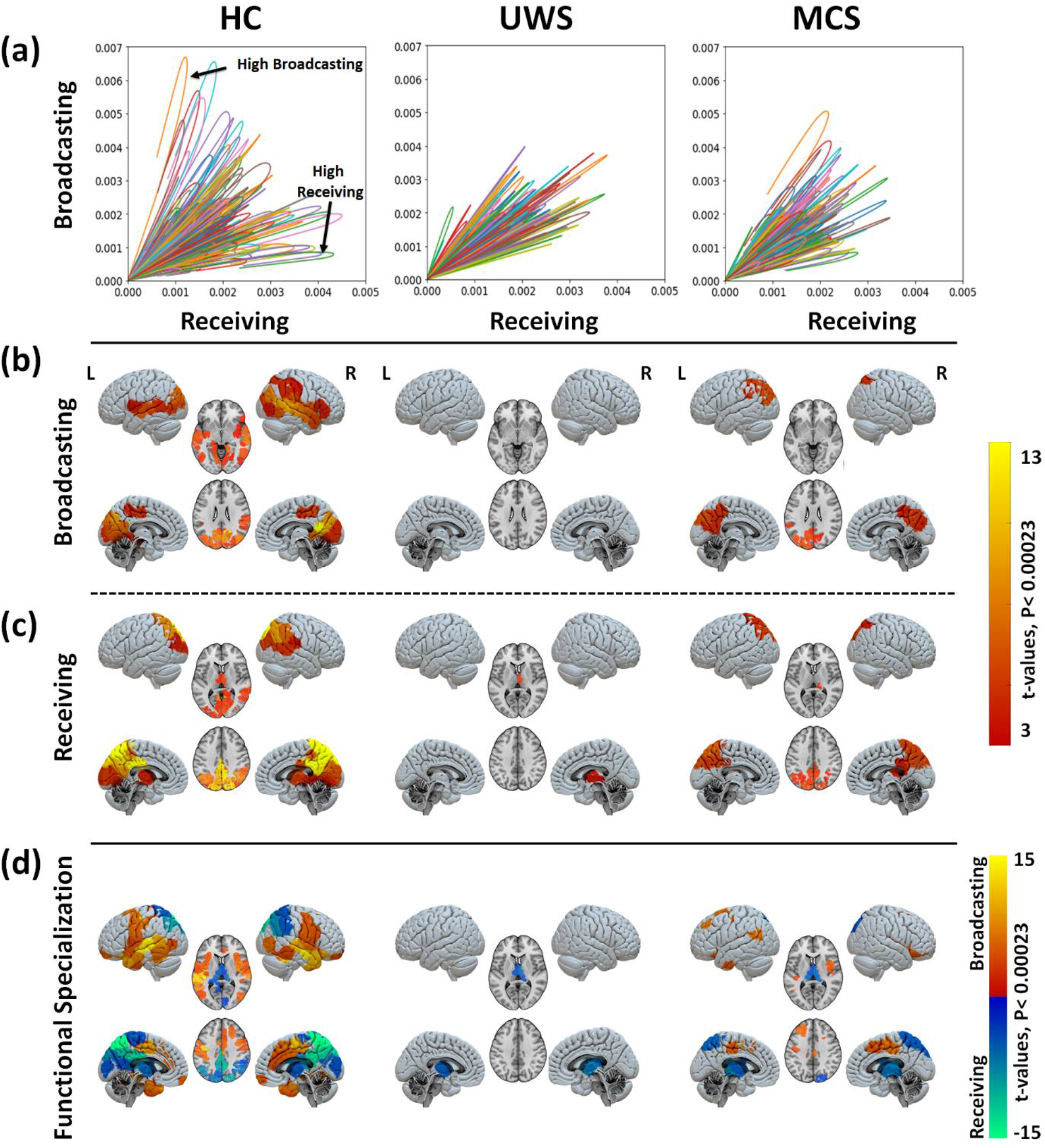
Region-wise broadcasting and receiving capacities due to exogenous perturbations. (a) Leveraging the asymmetric properties of the response matrices presented in a, the time course of the response can be plotted to represent the broadcasting and receiving properties per brain region (represented by differently colored lines). Healthy volunteers, and to a lesser extend MCS patients, are characterized by regions more dominantly involved in broadcasting or receiving. This functional organization is lost in UWS patients, who present an almost uniform distribution of regional properties with attenuated broadcasting and receiving. Please note that the end of the network response curve is cut at the end of the modelled time (200 s). The y and x axis is the network response for broadcasting and receiving capacities (unit free). Maps of significantly large broadcasting (b) and receiving capacities (c) for the three study groups (healthy controls, HC; unresponsive wakefulness syndrome, UWS; and minimally conscious state, MCS). The color code represents the t-values. Only regions with significantly high values are presented in each case (FDR corrected p-values <0.05 for 214 tests (ROIs)). (d) By subtracting the AUC of the **R**(t) curve for receiving from the AUC of the broadcasting curve in a region-wise fashion, we represent the dominant role of each region per group. From this it becomes clear that, although some regions play a role in broadcasting to and receiving from the network (e.g., the PCC area in MCS patients, or the occipital cortex and PCC in HC) the network most involved in receiving comprises the occipital and posterior cortical areas while broadcasting is the predominant role of the frontal cortices.

We further characterized the functional specialization of distinct regions by creating an anatomical map of the quantitative regional broadcasting and receiving properties. **Figure 4** reveals that the three levels of consciousness are characterized by distinct spatial distributions of *broadcaster* (**Fig. 4b**.) and *receiver* (i.e., integrator; **Fig. 4c**.) areas. Notably, in UWS patients no brain region stands out either as a broadcaster or as an integrator, except for the thalamus which displays a relatively large receiving capacity. In the healthy controls, we found several regions with both significant broadcasting and receiving capacity: the bilateral occipital, calcarine, lingual, cuneus, precuneus, superior and inferior parietal, right middle temporal. Significant broadcasting-only capacity was found in the bilateral middle and superior temporal, right inferior temporal, left parahippocampal, bilateral insula, inferior parietal, right supramarginal and right inferior frontal areas. On the other hand, the bilateral thalamus, posterior cingulate cortex (PCC), precuneus, middle cingulate and right supramarginal gyrus displayed significant receiving-only capacity. The MCS patient group was characterized by globally reduced broadcasting and receiving properties compared to healthy controls, however they showed a relatively preserved receiving and broadcasting of information within bilateral occipital, cuneus, bilateral superior and inferior parietal and precuneus. Additionally, they presented preserved broadcasting in left supramarginal and receiving capacity in the right thalamus, see **Fig. 4c. and Supplementary Table 3**. After quantification of the regional roles in broadcasting and receiving within the network, we evaluated the functional specialization of each brain area **Fig. 4d** by subtraction of the AUC of the **R**(t) curve for receiving from the AUC of the broadcasting curve in a region-wise fashion. This clearly shows two distinct networks, a receiving network prominently represented by the posterior and occipital cortex and the thalamus, and a broadcasting network encompassing parietal, temporal and frontal cortices.

We ended our perturbative analysis by comparing the region-wise group differences of the patients in respect to the healthy controls, which are shown in **Fig. 5**. Following the severely hampered broadcasting and integrating capacity in UWS, reduced information broadcasting in UWS patients as compared to healthy controls was especially notable at the bilateral hippocampus, parahippocampus, thalamus, caudate, amygdala, putamen, insula, inferior/middle temporal, temporal pole, right superior temporal, fusiform, lingual, calcarine, occipital, anterior cingulate, right inferior and middle frontal cortices. The notorious lack of broadcasting capacity of the subcortical regions evidences a reduced activity of the whole network. A profound reduction of the capacity to receive information in the UWS patients compared to the healthy controls was found at the bilateral precuneus, PCC, lingual, calcarine, fusiform, middle occipital, middle / anterior cingulum, inferior / superior parietal, supramarginal, middle temporal, inferior frontal cortices and the middle prefrontal cortex. These regions encompass primary visual and auditory areas, but also higher integration areas in the PCC that have an important hub function within the whole-brain network, see **Fig.5** **and Supplementary Table 4**.

**Figure 5.**
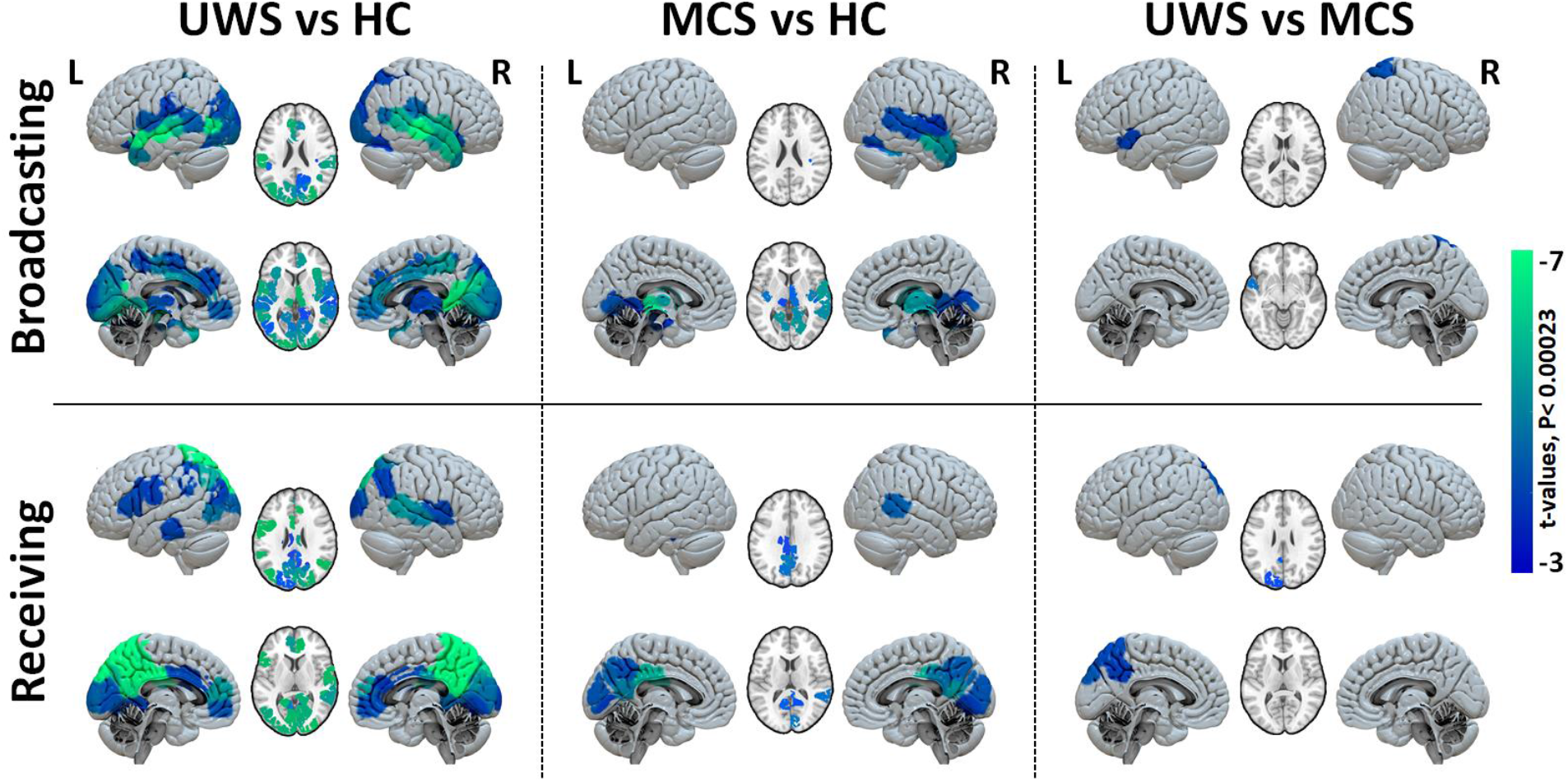
Group comparison of regional broadcasting and receiving capacities after exogenous perturbations. Maps of regional contrasts in broadcasting and receiving capacity between study groups. Color bar represents the t-values for regions with significant between-group differences (Bonferroni corrected p-values <0.05 for 214 tests (ROIs)).

The MCS patients showed a less pronounced picture of impaired information in- and out-flows. Compared to healthy controls they showed a significant reduction in the potential to broadcast information in the bilateral thalamus, parahippocampus, left hippocampus, bilateral insula, inferior / middle temporal, right superior temporal, bilateral fusiform and lingual cortices. A reduced capacity to receive information at the bilateral precuneus, PCC, cuneus, right lingual, calcarine, bilateral middle cingulum and right middle temporal cortices. Finally, compared to MCS patients, UWS patients showed additional significant reduction in receiving and broadcasting of information at the left precuneus, occipital cortex, temporal and right superior parietal, thus indicating that the information flow in these areas might be the most important contributors to conscious information processing.

### 3.6. Association of whole-brain network responses to clinical measures

Last, we explored how the computational measures employed here were associated with other aspects of clinical interest. We first evaluated if the whole brain responses (as in **Fig. 3a**) are sensitive to etiology alongside level of consciousness. Main and interaction effects of diagnosis and etiology on the peak value (i.e., early response) and late global brain responses (area-under-the-curve in the 60-200 sec of modeled time) were quantified using a linear regression model. We did not find significant differences for peak global responses for diagnosis (p= 0.405), etiology (p= 0.137), or the interaction between diagnosis and etiology (p= 0.258), the model r-squared = 0.27 and model p-value = 0.008. For the late global response, we noted significant differences in the case of diagnosis but neither in etiology nor in the interaction between diagnosis and etiology (p-value for diagnosis = 0.0004, p-value for etiology = 0.063, p-value for interaction = 0.096; the model r-squared = 0.32 and model p-value = 0.0027). This confirms that the main differences in our analyses are due to diagnosis and ensured that the observed effects were mediated by the level of consciousness primarily, rather than the different neurobiological effects of etiology. Indeed, looking at the global brain responses of individual subjects it shows that the anoxia and TBI subjects are diversely distributed in both UWS and MCS patient group (**Fig. 6a**). As etiology and the interaction between etiology and diagnosis were not significant, we did not explore their relationship with the (regional) communicability responses further.

**Figure 6.**
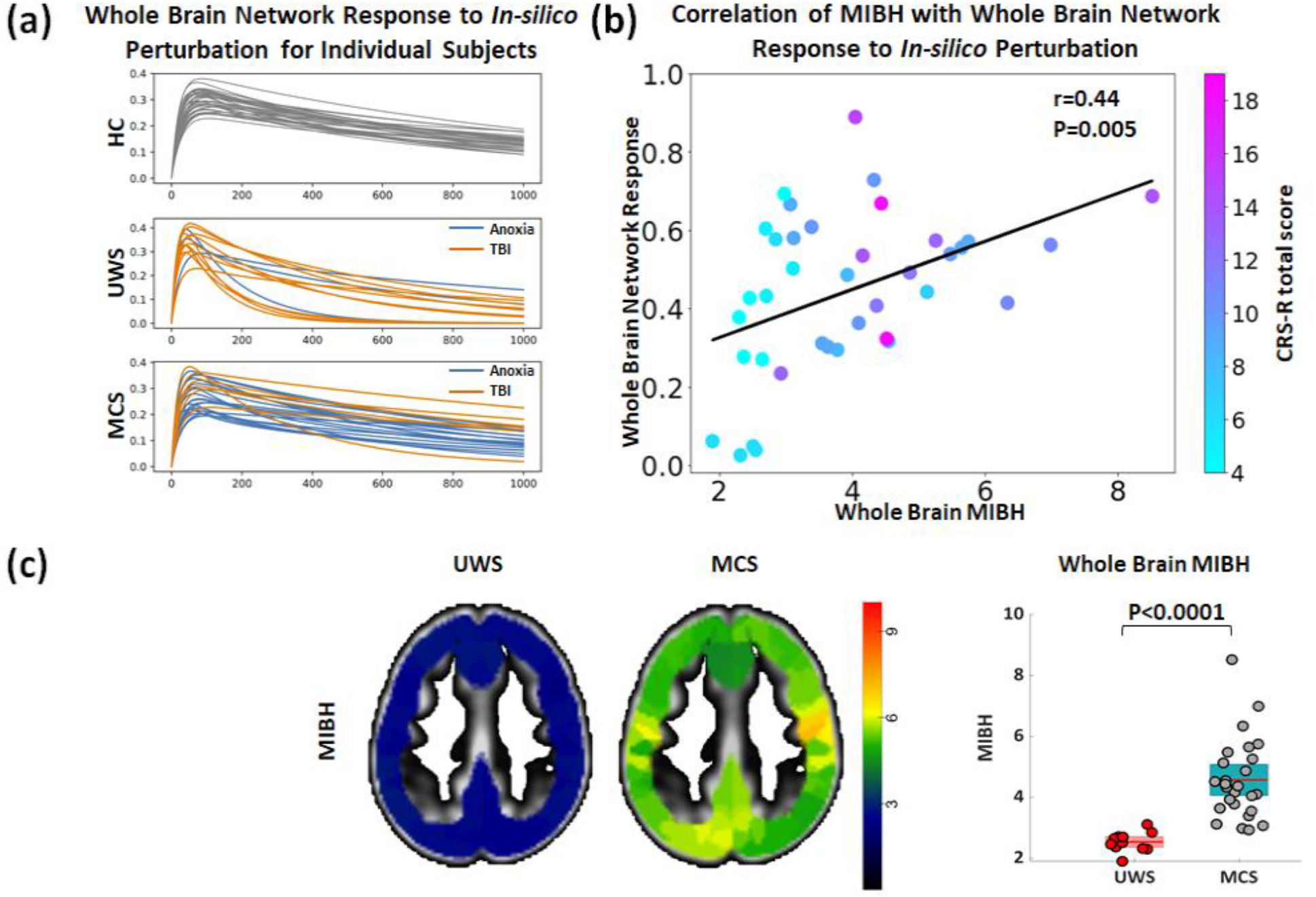
Association of the individual whole brain response curves of *In-silico* perturbations and clinical variables. (a) **R(t)** Response curves are grouped by diagnostic entity including healthy controls (HC; top), unresponsive wakefulness syndrome (UWS; middle) and minimally conscious state (MCS; bottom) and colored by etiology in the case of DoC patients (Anoxia in orange and non-Anoxia (e.g., TBI, hemorrhage) in blue). Anoxia and TBI patients whole brain network response are heterogeneously distributed in both UWS and MCS groups. (b) Correlation of whole brain response to *In-silico* perturbation (y-axis) with the metabolic index of best hemisphere (MIBH; x-axis). Individual datapoints are color coded according to the patient’s CRS-R total score. A positive and linear relationship between the network response and glucose metabolism is present, that is also associated to the presence of more conscious behaviors (CRS-R total score). (c) Shows an example for the MIBH images for a representative UWS and MCS patient. As also evident from the boxplot, glucose metabolism is minimal in UWS patients while partially preserved in MCS patients.

Since information processing requires the consumption of energy at the neural level, we quantified the neurobiological underpinnings of the integrative network response by establishing a direct link between cerebral glucose metabolism (as measures with glucose PET imaging) and network integration (as measured with the whole-brain computational response to perturbation). Specifically, we correlated the whole brain network response of the regions that showed different network responses in HC and UWS patients with glucose PET MIBH. The MIBH of UWS was significantly lower (2.53±0.31) as compared to MCS patients (4.57±1.32)., p<0.0001) (**Fig. 6c**). The association of the complex model-based assessment of network function to basic neural function in the first place helps to increase confidence in the model. Second, although a causal relation cannot be established to date, it is a first step towards the exploration of the mechanism behind the widespread observed glucose metabolic changed in DoC patients. We noted positive correlations of MIBH with the whole brain network responses (r=0.44, p=0.005) (see **Fig. 6b**). Indeed, patients with more preserved MIBH and functional network responses show more complex behaviors as evidenced by the clinical evaluation with the CRS-R (see color coding in **Fig.6b.**).

In summary, our model-based analyses to estimate effective connectivity and to simulate *in-silico* the response to exogenous perturbations can be related to basic biological measures and clinical observations, allowing us to identify specific directed pathways that are disrupted in patients with DoC.

## 4. Discussion

In the present paper, we have studied the neural propagation of endogenous and exogenous perturbations in the brain using model-free and model-based analysis methods, applied to the problem of elucidating the mechanisms behind the loss of consciousness due to acquired brain injury and its partial recovery. The methods here employed add significant value to the dynamical approaches for two main reasons. First, they rely on simple observables – the resting-state fMRI – and thus they do not require the execution of experimental exogenous stimulation protocols. And second, unlike previous approaches, they allowed us to investigate the directional causal interactions between brain regions, thus elucidating alterations in the broadcasting and the integrating capacities of individual areas or pathways, between normal awake and unconscious patients. Our main finding is that we could identify two distinct malfunctioning neural circuits in patients with DoC: the posterior cortical regions fail to convey information, in conjunction with reduced broadcasting of information from subcortical, temporal, parietal and frontal regions. These results show that patients with prolonged disorders of consciousness lack of the capacity for the integration of events that would lead to conscious perception.

In healthy controls we found that the relaxation time constants associated with the resting-state BOLD signals display a gradient distribution with shorter relaxation times in subcortical areas and longer time constants in the frontal and in the parietal areas, **Supplementary Fig. 2a**. Accordingly, analysis of exogenous *in-silico* perturbations revealed that the broadcasting of information flow is predominant in a broad range of cognitive modules, including the hippocampus, parahippocampal, temporal, posterior and inferior frontal regions. This subcortical-cortical loop has been proposed to mediate the sensory information to be globally ‘accessible’ to other cognitive functions through feedforward and feedback loops by the global neuronal workspace theory, and only when access to all cognitive modules occurs, sensory content is elevated to conscious perception (Dehaene et al., 2011; Dehaene and Changeux, 2011; Mashour et al., 2020). Although the activity in the posterior regions is highly influenced by perturbations, suggesting that they have a large cause-effect capacity to receive and integrate the information flow, one of the key principles of conscious perception in integration information theory (Oizumi et al., 2014; Tononi, 2004; Tononi et al., 2016).

Regarding the patients with unresponsive wakefulness syndrome (UWS), the results observed were very much altered. First, the propagation of endogenous events occurring in the resting-state BOLD rapidly decay avoiding their subsequent integration, **Fig. 1a**. This is corroborated by the fact that the spatial distribution of relaxation times fades away in the UWS patients, with all areas taking short relaxation times (**Figs. 1b** and **c**) and evidencing that local activity does not properly propagate along the network. Especially frontal, parietal and higher-order cortices which ensure sensory information processing, need longer time to integrate diverse information (Hasson et al., 2008; Yeshurun et al., 2017). Second, effective connectivity is reduced overall but interestingly some effective subcortical-cortical connections were found to significantly increase, **Fig. 2b**. The propagation of *in-silico* exogenous perturbations showed a rapid and large response followed by a fast decay, **Fig. 3**, since the information fails to propagate along the network in a sustained manner. Such early hyper-response has been previously seen in UWS patients (Di Perri et al., 2013) and in loss of consciousness due to generalized epilepsy possibly caused by an excess of electrical discharges in the brain (Moeller et al., 2008). The mechanism for this hyper-response, **Supplementary Fig. 3**, is yet to be elucidated but we envision two possibilities: it could arise either due to the network being dominated by short local recurrent loops, or due to a lack of inhibition as in unconscious anesthetized ferrets (Wollstadt et al., 2017). Finally, the regional specificities for broadcasting and receiving that were observed in the healthy controls are vanished for the UWS patients. Only the thalamus stands-out, as an area with significant receiving capacity thus probably allowing its gateway function between the body and the brain.

The patients in minimally conscious state (MCS) studied here underwent through a coma and a UWS phases after brain injury, but later regained partial consciousness and cognitive functionalities. All the results found for the MCS patients show light alterations to those in the healthy participants, as expected from their partial functional recovery. The spatial gradient of relaxation time constants from the resting-state BOLD is recovered, **Supplementary Fig. 2**, except shorter time constants were still found in the thalamus and the left medial prefrontal cortex. Effective connectivity was in general slightly below values observed in control subjects but a significant reduction was found remaining in fronto-parietal connections and in between temporal regions, **Fig. 2c**. Regional broadcasting and receiving capacities to *in-silico* perturbations displayed a recovered scenario, see **Fig. 4** and see **Supplementary Table 3**. It is clinically relevant to understand why or how MCS patients could partially recover from the unresponsive wakefulness state. In the light of our results, it seems that a sufficient regain of the propagation of information leading to an increase in receiving and broadcasting of information at the left precuneus, occipital cortex, temporal and right superior parietal is instrumental for the recovery of conscious information processing, **Fig. 5**.

In comparison with healthy controls, the UWS patients showed a reduction of receiving information in posterior regions, which implies sensory information integration is impaired already at the level of sensory regions to the high-level hub regions of the Default Mode Network (i.e. PCC and Precuneus). Indeed, the lack of receiving of information in the sensory areas hampers a stimulus or event to reach awareness, as integration of external inputs is a prerequisite for consciousness (Herbet et al., 2014). Our results provide a mechanistic explanation for how the ability to receive information in sensory and DMN hub regions alters cerebral information processing, which might be at the essence for the structural and functional anomalies in UWS patients. Although it is known that the PCC and Precuneus have decreased structural, functional and metabolic integrity, (Annen et al., 2016; Demertzi et al., 2015; López-González et al., 2021; Luppi et al., 2019; Stender et al., 2014) our results show that the ability to receive information in sensory and DMN hub regions is reduced in UWS patients. On the other hand, broadcasting was reduced in the subcortical regions (i.e., thalamus, caudate, putamen) and regions involved in higher cognitive function (i.e., temporo-parietal, anterior cingulate and frontal regions). This is aligned with the mesocircuit hypothesis (Schiff, 2010) which states that the feedforward connections between these regions play a key role in reaching levels of (cortical) activity that support the stimuli to access consciousness processing. This has been recently confirmed in macaques using intracraneal-EEG, showing that integration at the thalamus, caudate, putamen and parietal cortex is a hallmark of conscious states (Afrasiabi et al., 2021). Our findings unravel that human consciousness also relies on the broadcasting capacities (i.e., from the thalamus, caudate and putamen) to support the transmission of activity, functional integration and recurrent activity between subcortical and cortical neurons, all of which is lost in UWS patients. Interestingly, we also noted a decrease in receiving and broadcasting capacities in the bilateral temporal areas for DoC patients. Recent studies also noted altered structural (Annen et al., 2018) and functional loss in the temporal area in DoC patients (Demertzi et al., 2015; Thibaut et al., 2021). To date, there is limited explanation for the involvement of the temporal cortex in consciousness. We speculate that a lesser involvement of the temporal areas in the information pathways could impede self-awareness and memory.

From a clinical point of view, there is a variety of causes leading to disorders of consciousness. These are typically classified into two categories. On the one hand, *anoxia* is a generalized damage of brain tissue due to a temporary disruption of oxygen supply caused by, e.g., heart attack or asphyxia. On the other hand, loss of consciousness can also occur due to focal brain lesions due to, e.g., traumatic brain injury, stroke or epilepsy. It is well-known that the probability of (partial) recovery from UWS is larger for patients with focal lesions than for anoxic patients (Thibaut et al., 2021). This is reflected in the sample of patients here studied since 11 out of 14 patients with UWS are anoxic and 19 out of 26 patients in MCS suffered from focal brain injuries. However, the extent of damage is not a perfect predictor and the heterogeneity of lesions across patients calls for a better understanding of the mechanistic causes leading to the loss and the (partial) recovery of consciousness. We have explored the effects of etiology in our results and found that the diagnostic classification of the patients into UWS and MCS based on EC and on the propagation of perturbations significantly correlate, while this was not the case for etiology. Even though at this point we could not establish a link between etiology and network responses, perhaps due to a lack of data and heterogeneity within the data, hypothetically the evaluation of lesion-specific alterations in network responses could help formulate more specific predictions about where propagation is altered in reduced conscious states. Other samples including healthy volunteers under sedation could help narrowing down the location of possible consciousness mechanisms further as well.

Additionally, we have observed the presence of information processing leading to consciousness follows relatively widespread preserved glucose metabolism (**Fig. 6**). This potential neurobiological basis for information processing is associated to the richness of signs of consciousness at the behavioral level as well. However, the causal relationship between regional glucose metabolism and their broadcasting / receiving properties is still to be clarified. On the one hand, we could speculate that a significant disruption of the propagation of events along the neural network leads to a loss of metabolic demand, which is then observed via glucose PET. But, on the other hand, the opposite possibility also needs to be considered that the brain lesions could disrupt the metabolic supply chains, thus exhausting the capacity of the neural network to generate sufficient activity and responses.

Our *in-silico* pertubational study revealed transient global changes to the dissemination of information, associated to glucose metabolism, that are similar to those we observed from the integration of endogenous events with the intrinsic ignition. In DoC patients, and especially in UWS patients, the brain’s cause-effect capacity to respond is significantly lower than during normal wakefulness in healthy subjects. It seems that the observed spatiotemporal alterations of local event processing also hamper global integration and whole brain neural responses; as observed both after *in-silico* and endogenous perturbations. These results are in line with empirical studies using TMS, in which the recruitment of global neural activity after perturbation, both in space and time, has been found to be reduced during deep sleep, anesthesia and in DoC (Casali et al., 2013; Massimini et al., 2005). In conclusion, the cerebral capacity of propagation and integration of local, naturally occurring events into the entire network is affected by reduced states of consciousness and shares similarities with both information integration theory (Tononi, 2004; Tononi et al., 2016) and global neuronal workspace theory (Dehaene et al., 2011; Mashour et al., 2020). Although these theories have distinct concepts of consciousness, our results suggest that they might represent two sides of the same coin (Northoff et al., 2020; Northoff and Lamme, 2020; Winters, 2020).

## Abbreviations

BOLD: Blood oxygenation level dependent
[^18^F]FDG-PET: Fluoro-deoxyglucose Positron Emission Tomography or glucose PET
fMRI: functional MRI
CRS-R: Coma Recovery Scale-Revised
DoC: Disorders of consciousness
HC: Healthy control
MCS: Minimally conscious state
UWS: Unresponsive wakefulness syndrome
PCC: Posterior cingulate cortex
SC: structural connectivity
EC: effective connectivity
τ: Relaxation time constants
MOU: multivariate Ornstein-Uhlenbeck

## Acknowledgements

We would like to thank the healthy participants and the patients, their families, caregivers and treating clinicians for their participation in this study. The authors thank the whole staff from the ICU and Nuclear Medicine departments, University Hospital of Liege. We are highly grateful to the members of the Liege Coma Science Group for their assistance in clinical evaluations.

## Authors’ contributions

RP, ALG, JA, GZL, MG, GD and SL designed research. JA, SL and GZL supervised the research. JA, AT, CM, OG and the Coma Science Group Collaborators acquired the data. RP, ALG and AE preprocessed the data. RP, ALG, JA, GZL and GF analyzed the data. MG and GZL designed the computational model and optimized the code as per this research study. RP, JA, ALG and GZL wrote the manuscript. All authors interpreted the results and contributed to the editing of the manuscript.

## Collaborators

Alice Barra, Charlotte Martial, Claire Bernard, Estelle Bonin, Emilie Szymkowicz, Jean-Flory Luaba Tshibanda, Leandro Sanz, Marie Vitello, Roland Hustinx.

## Figure legends

**Supplementary Figure 1.**
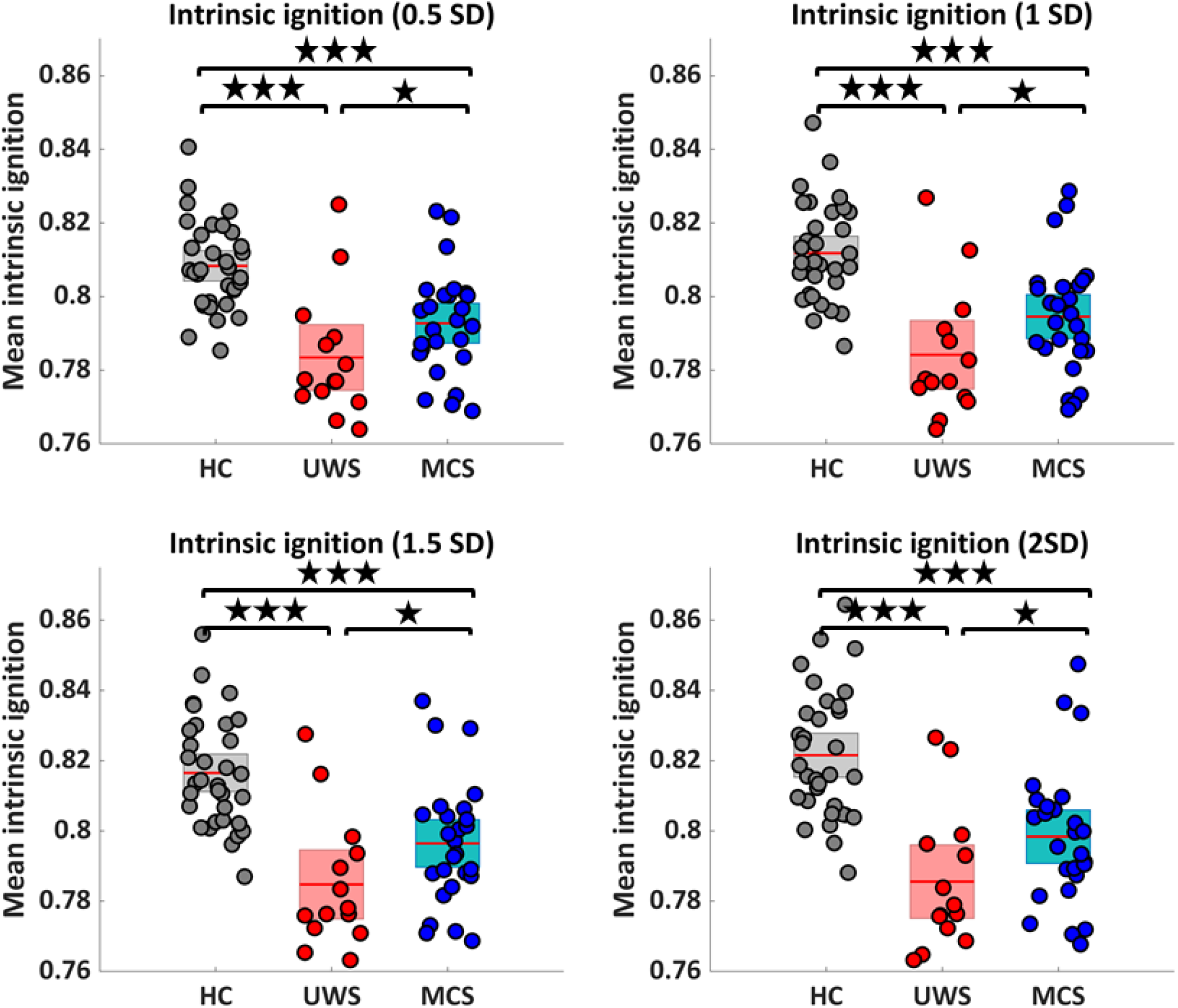
Comparison of different SDs for the event threshold in the BOLD time series to detect intrinsic ignition events. Mean intrinsic ignition in healthy controls (HC) unresponsive wakefulness syndrome (UWS) and minimally conscious state (MCS) are presented. As the event threshold may affect integration values, we used 0.5 to 2SD event threshold to compute the intense ignition. However, we found similar patterns for all the thresholds (i.e., 0.5 to 2 SD), illustrating the reduced capacity to integrate endogenous spontaneous events in patients with DoC. Please note that intrinsic ignition could not be computed for higher thresholds due to lack of events. Stars reflect the Bonferroni corrected (for three groups) significance levels (* = p <0.016; ** = p <0.001; *** = p <0.0001).

**Supplementary Figure 2.**
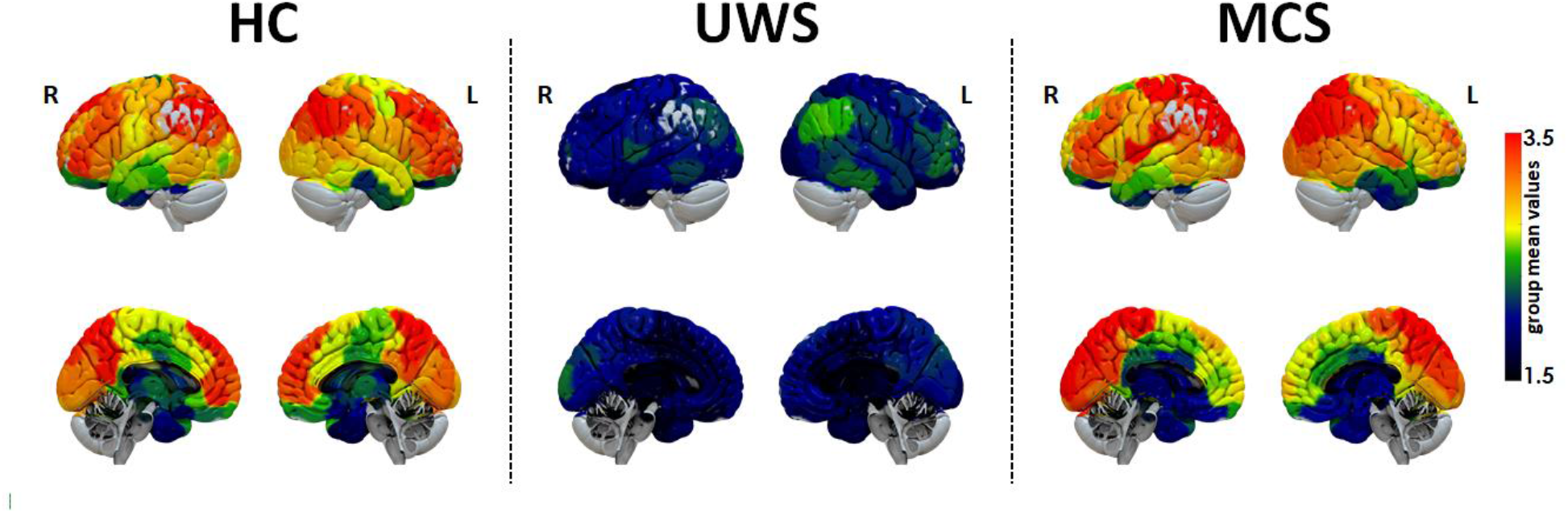
Spatial maps showing the regional distribution of relaxation time constants (τ) as calculated for empirical BOLD signals for each area. Healthy controls display a spatial heterogeneous distribution of τ, while UWS patients are characterized by short time constants overall. The spatial distribution of regional τ is very much recovered in MCS patients.

**Supplementary Figure 3.**
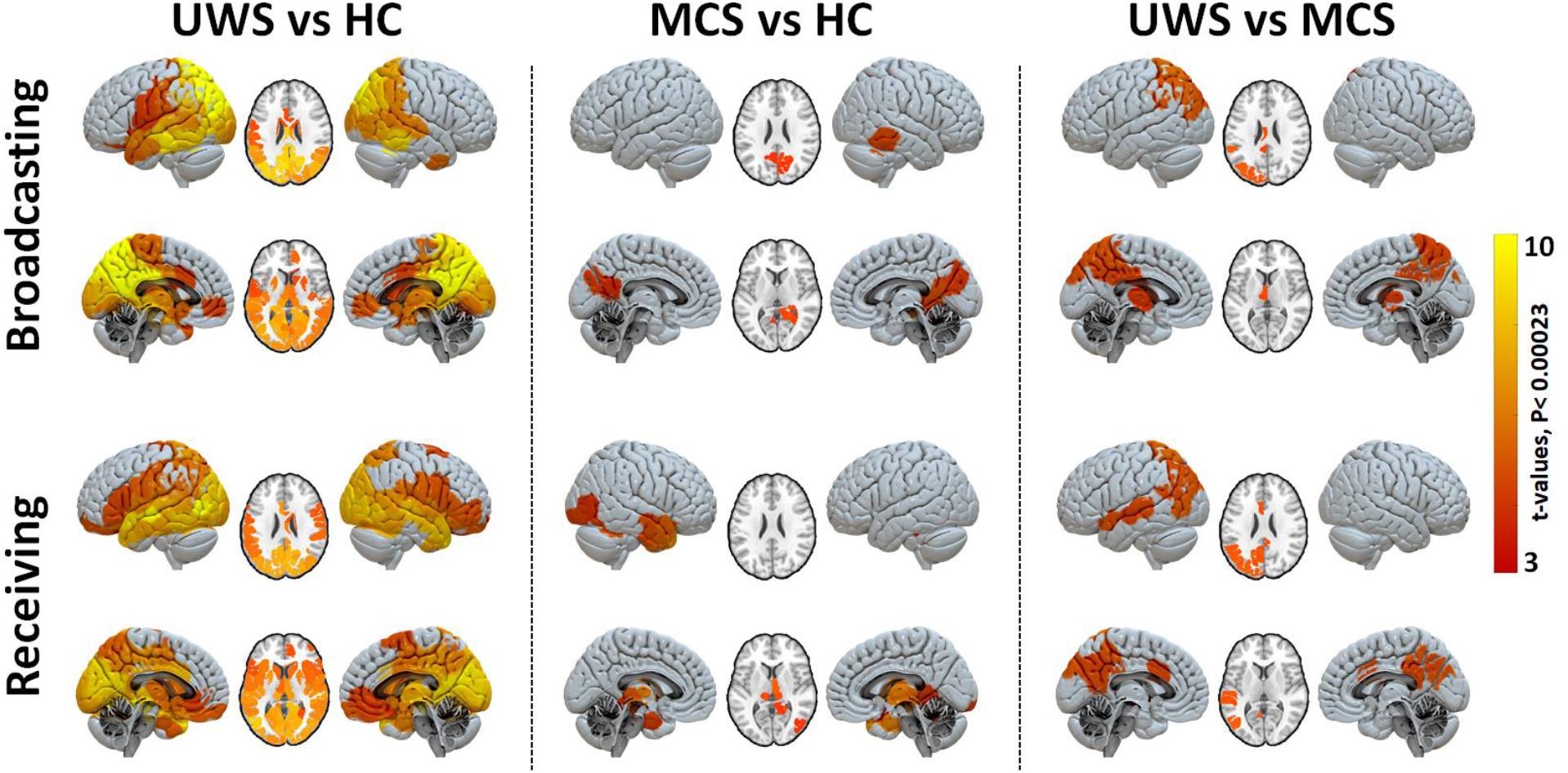
Brain regions' hyper-response (higher growth with sudden decay) differ in (a) UWS patients compared to HC (b) MCS patients compared to HC (c) UWS compared to MCS patients for receiving and broadcasting capacities. Color bar represents the t-values for regions with significant between-group differences (Bonferroni corrected for 214 tests).

